# IsoSense: Frequency Enhanced Sensorless Adaptive Optics Through Structured Illumination

**DOI:** 10.1101/448613

**Authors:** Mantas Žurauskas, Ian M. Dobbie, Richard M. Parton, Mick A. Phillips, Antonia Göhler, Ilan Davis, Martin J. Booth

## Abstract

We present IsoSense, a wavefront sensing method that mitigates sample dependency in image based sensorless adaptive optics applications in microscopy. Our method employs structured illumination to create additional, high spatial frequencies in the image through custom illumination patterns. This improves the reliability of image quality metric calculations and enables sensorless wavefront measurement even in samples with sparse spatial frequency content. We demonstrate the feasibility of IsoSense for aberration correction in a deformable mirror based structured illumination superresolution fluorescence microscope.

## 1 Introduction

Sensorless adaptive optics [1] is an aberration measurement technique, in which optimum aberration correction is inferred from a collection of intentionally aberrated images. It has found widespread use in various forms of high resolution microscopy, in addition to optical coherence tomography[2, 3], spectroscopy [4] and laser communications [5]. In microscopy, sensorless AO is particularly useful for correction of specimen induced aberrations when other, sensor-based, methods are not feasible. It can be used as a standalone wavefront sensing method or in conjunction with sensor-based techniques [6].

Sensorless AO works by applying different amounts of bias for chosen aberration modes and assessing the image quality by calculating a pre-selected quality metric for each image. Depending upon the application, the metric could be based upon, for example, intensity, contrast or the spatial frequency content of the image. The necessary aberration correction is then estimated from the metric measurements using a mathematical model of the imaging process. The main challenge here is that the quality metric must be chosen carefully to work well with particular classes of imaged objects [7]. However, even when the metric is chosen using prior knowledge about the sample, the measurements still rely on the local structure of the object that was imaged. In microscopy, sensorless AO techniques commonly rely on two groups of image quality indicators. First is signal intensity [8, 9]; the second relies on structural information, for example spatial frequency content optimization (here, the term “Spatial frequency content” refers to the amplitude of sinusoidal components, as determined by Fourier analysis of an image.) [10, 11, 12] or wavelet decomposition [13]. In some limited cases, phase retrieval has been successfully demonstrated [14, 15, 16], but it is only applicable to a narrow range of specimens.

Previous work has observed that the performance of the image-based algorithms are, to some degree, dependent on the specimen structures present. When the specimen contains a fairly isotropic distribution of structures – or, more specifically, spatial frequencies – then the algorithms behave reliably. Conversely, when there is a strong anisotropy to the structures, such as when the specimen consists of aligned linear components, there can be a bias in the measurements. The reason for this bias is that certain aberration modes have a greater effect than others on particular spatial frequencies. It follows that the modes to which the system is insensitive are not effectively corrected [17].

Here we propose a sensorless AO method which relies on structured illumination to create multiple frequency-shifted copies of image information to ensure adequate sampling of the microscope optical transfer function (OTF). Our method improves the sensitivity and reliability of sensorless AO when used for imaging typical objects: it permits accurate assessment of image quality and enables precise correction of sample-induced aberrations to recover the optimal OTF of the instrument for widefield fluorescence imaging. We have demonstrated its performance in structured illumination microscopy (SIM) [18, 19, 20], because this technique is particularly sensitive to the fidelity of imaging high spatial frequencies [21], and the equipment for its implementation is readily capable of forming custom illumination patterns.

## 2 Principles of IsoSense

### 2.1 Image quality based wavefront sensing

Image quality based wavefront sensing relies on using images obtained with the microscope as an input for an algorithm that estimates the optical aberrations present in the system. A classical implementation of sensorless AO is depicted in the flowchart (Fig. 1a). Here, a series of object images are acquired with different amounts of selected aberration modes applied with an adaptive optical element. Ideally, the aberration modes form a set of orthogonal eigenmodes so that each has an effect on the image quality that is independent of the other modes. In practice, a standard basis set, such as Zernike polynomials, is used for convenience. An image quality metric is selected and evaluated for each image (Fig. 1b). A parabola (or other suitable function) is fitted to the measured points and the mode coefficient corresponding to the estimated peak is applied as the correction. Subsequent modes are then corrected in a similar manner (Fig. 1c). For the example presented in our paper we were sequentially correcting pre-selected Zernike modes to reach the optimal correction. In particular circumstances, it has been shown that to correct n aberration modes, a minimum of 2*n* + 1 measurements are needed. However, more measurements may be beneficial where the signal-to-noise-ratio (SNR) of the data is low.

**Figure 1:**
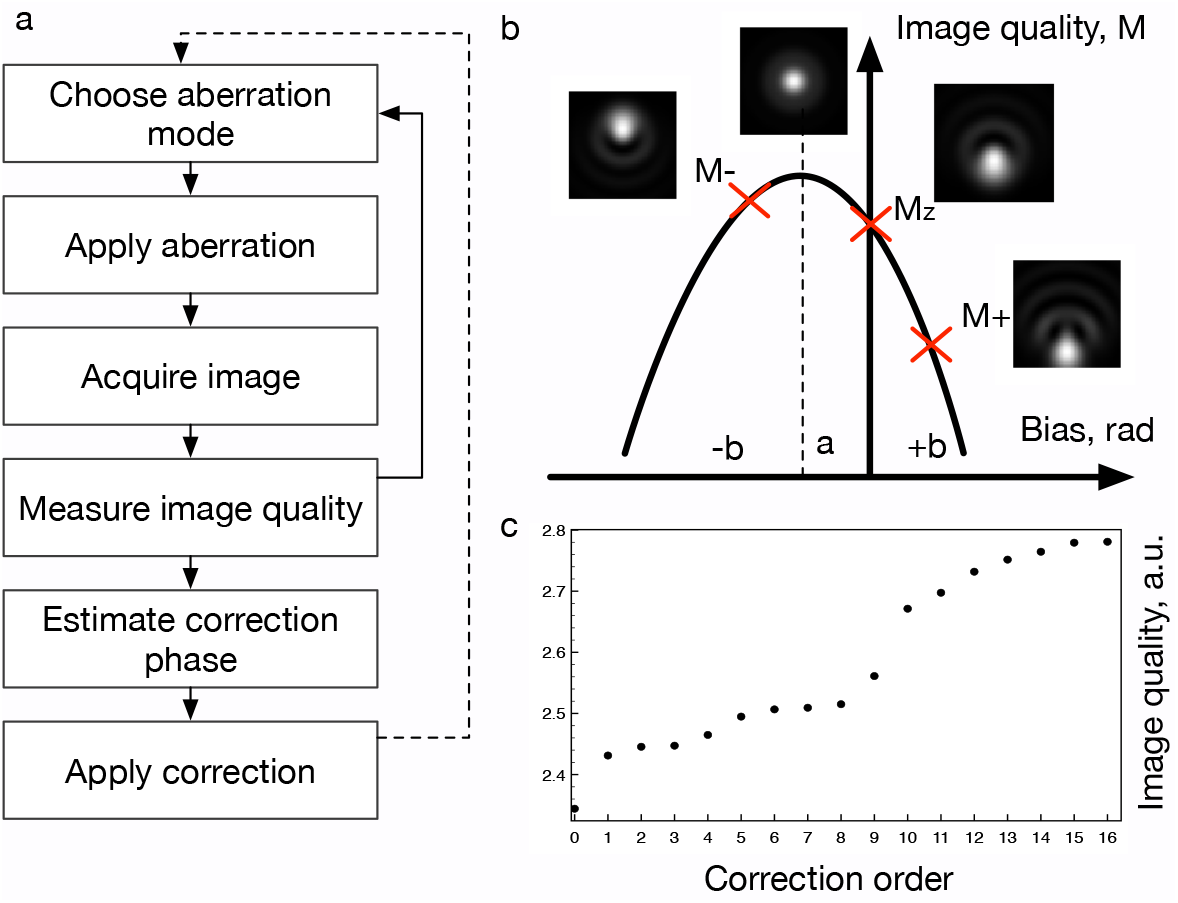
Image quality metric based wavefront sensing. (a) Flowchart depicting classical implementation of sensorless AO based wavefront measurement. (b) Sample images are captured for each mode using at least three different bias values (-b, a, +b). Image quality is estimated using a suitable defined metric for each image (M-, Mz, M+) and a quadratic function is fitted to the measured points. The peak value of the fitted curve corresponds to estimated best correction. Inset images represent a point spread function affected by various amounts of coma. (c) Typical image quality evolution as different modes are corrected in pre-set sequence.

Ideally, the image quality metric chosen for an imaging system should be sensitive to small changes in aberrations and would be compatible with a wide variety of samples. Typical characteristics associated with high-quality fluorescence microscopy images are brightness and sharpness. Optical aberrations reduce high spatial frequency content in widefield fluorescence images, reducing image sharpness and also affect the light intensity distribution through broadening of the point spread function (PSF). Quality metrics that exploit intensity distribution or brightness tend to be either insensitive in widefield mode (as in the case of total brightness measurements) or dependent upon object structures (as observed with histogram based or contrast measurements); they can also be affected by bleaching. Metrics that rely on sharpness are more robust against bleaching, but they also tend to be sample specific, working well only where the sample exhibits well-defined structures. In this paper, to numerically assess the image quality and calculate the quality metric, we used high spatial frequency content as a proxy for image quality. The merit function was calculated by using the absolute value of the image FFT, integrated over an annular region of the image spectrum from 0.15*R* to 0.85*R* where *R* was the radius of the calculated region of support of the image spectrum. We have omitted the central 15% of the information around the DC component to minimize the effect related to bleaching, and also we have omitted the outer 15% due to very low SNR in this region.

The OTF provides a way of understanding the operation of an imaging system in terms of its transmission of spatial frequencies. One does not normally have access to a measurement of the OTF, although the image spatial frequency spectrum may provide a proxy, if a wide spread of spatial frequencies is present in the object. A typical example of a sample that contains a wide range of spatial frequencies is a lawn beads with diameters smaller than the diffraction limit (Fig. 2 a, b, c, d). Correcting for aberrations shows an increase in the high spatial frequency content of the image.

**Figure 2:**
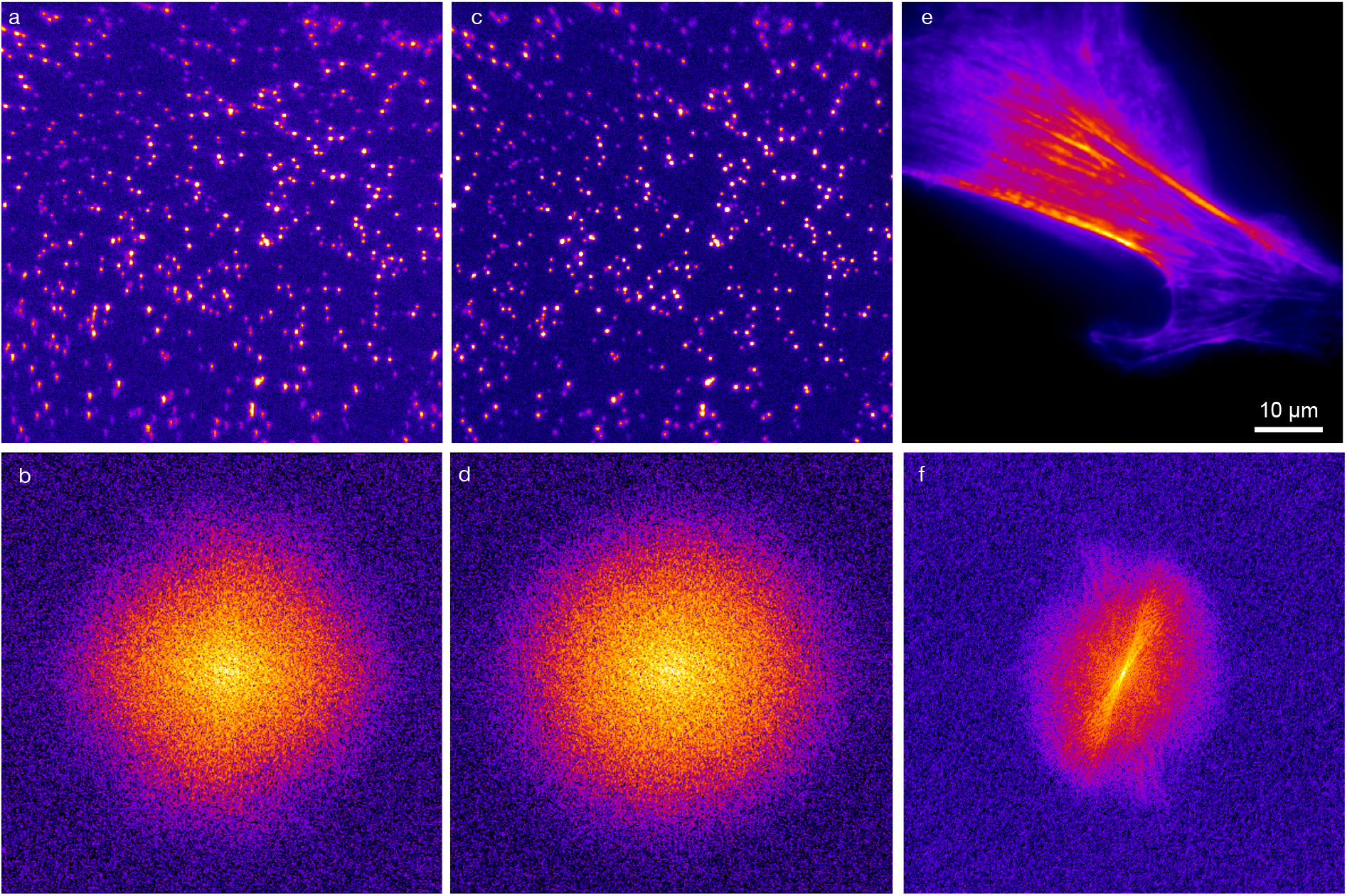
Spatial frequency content as a basis for image quality measurements. An FFT of a lawn of sub diffraction limit sized beads contains a good representation of all the spatial frequencies that can be transmitted by the system (raw image a, and FFT b) before correction and (c,d) after. In many experiments, images of real samples do not contain a wide enough range of spatial frequency information for adequate pupil sampling: an image of a bovine pulmonary artery endothelial cell showing preferentially oriented actin (e) illustrates uneven information distribution in the spatial frequency space (FFT) (f).

The figure 2e is an image of a bovine pulmonary artery endothelial cell imaged in the green (Alexa Fluor 488 phalloidin, labelling actin), while 2f is the amplitude component of its Fourier transform. The main structures in the image are preferentially oriented long actin filaments of about 7 nm diameter. This anisotropy in the sample means that the image spectrum does not sample the OTF uniformly: there is detailed information for some regions of the OTF, but little usable information for other regions. It is only possible to measure the effects of aberrations that affect the regions in the OTF for which information with a good SNR is present in the image spectrum. With some commonly occurring biological samples the signal can fall below the noise floor at mid or high spatial frequencies, resulting in sub-optimal adaptive correction. Therefore, an alternative approach is needed that can be applied to a wider variety of objects.

### 2.2 Custom illumination patterns for enhanced sampling of the OTF

In the simple case of a widefield, incoherent imaging system, the optical transfer function (Fig. 3a) can be calculated as the autocorrelation of the pupil function (Eq. 1) (Fig. 3b). Therefore, every point on the horizontal axis in Fig. 3b would correspond to a different magnitude of 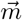 in Fig. 3a. The absolute value of OTF is numerically identical to the contrast of the image of a sine grating at the corresponding spatial frequency. An important consequence of this is obvious: the value of the optical transfer function at a particular value of 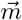 (or spatial frequency) is only dependent on the regions of the pupil function that have contributed to the calculation. In the (Eq. 1) ⋆ represent correlation and P* is a complex conjugate of the pupil function.

**Figure 3:**
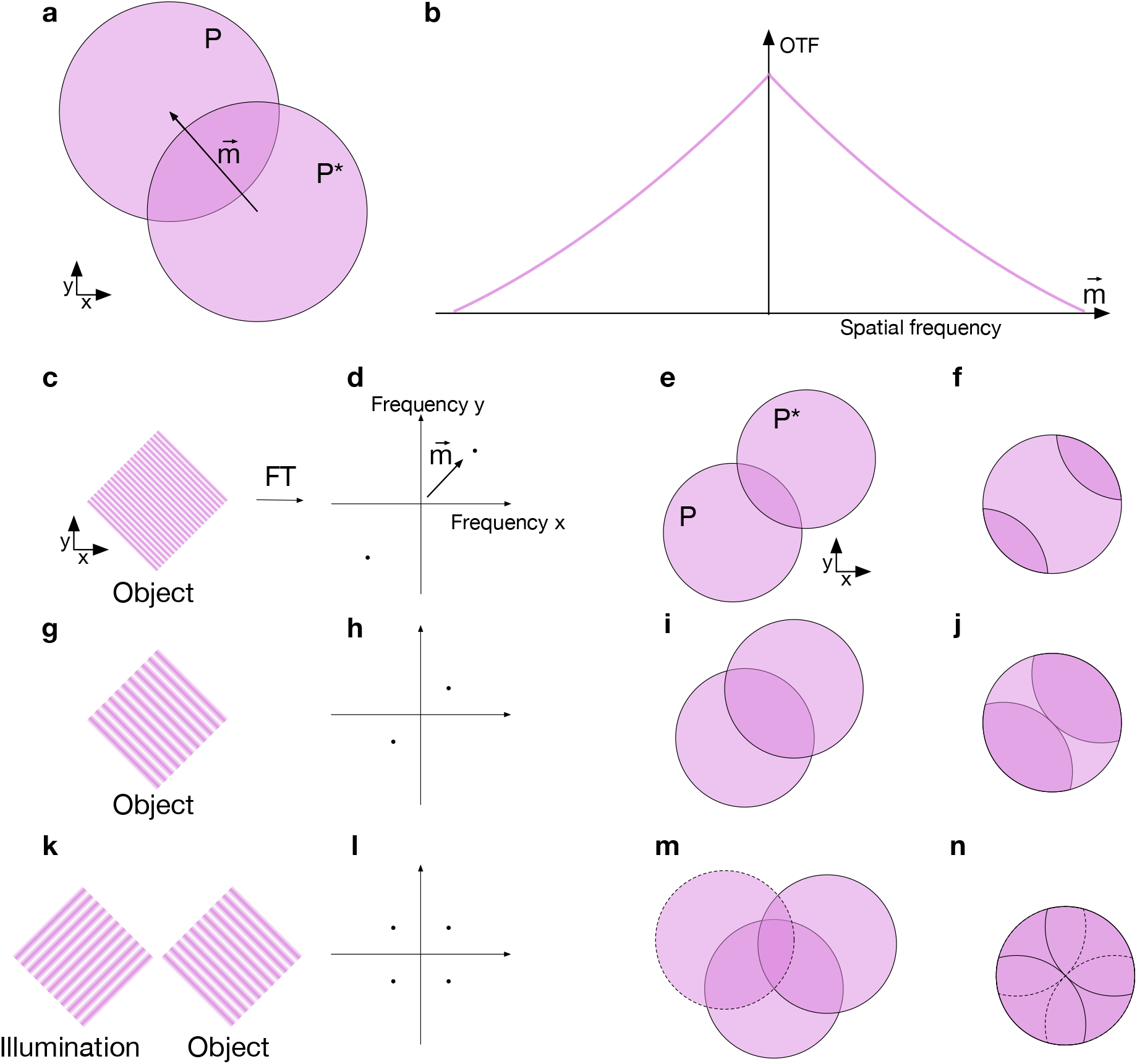
Custom illumination patterns for enhanced sampling of the OTF. (a,b) Illustration of OTF calculation. The OTF (b) is calculated as the auto-correlation of the shifted pupil function (a). The absolute value of the OTF is numerically identical to the contrast of the image of a sine grating at the corresponding spatial frequency. In a simplified case, an object consisting of one spatial frequency (a,b), would be affected only by aberrations affecting the OTF (e,i) through a few limited zones in pupil plane (f,j). Conversely, any phase aberrations that do not affect the spatial frequencies present in an object (denoted by dark pink colour, (f,j,n)) are undetectable by using image quality based metrics. (k,l,m,n) illustrates that a carefully chosen illumination pattern can help to fill the pupil with information and ensure adequate sampling of the OTF.

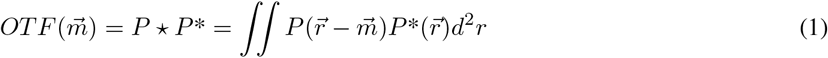

Figure 3 further illustrates the simplified case of an object which is defined by a single sinusoidal frequency; the object spectrum consists of a zero frequency component in addition to the positive and negative frequencies of the sinusoid. If we consider the contribution of the sinusoidal component, when the spatial frequency is high (Fig. 3 c, d, e, f), only the two dark-shaded segments of the pupil contribute to the imaging of this particular frequency. For lower frequencies, the information is distributed across larger areas of the pupil (Fig. 3 g, h, i, j). One consequence of this is that the quality of an image for an object containing only a limited set of spatial frequencies, the image quality is affected predominantly by aberrations in only a sub-set of the pupil function. The sensing method would be mostly insensitive to aberrations in the other pupil areas. IsoSense aims to eliminate these “blind spots” by applying structured illumination (Fig. 3 k, l, m, n) to fill the whole pupil with information.

It also should be considered that the efficiency of frequency transfer drops severely for high spatial frequencies (Fig. 3b) and OTF only can be measured if the information is above the detection noise floor. Therefore, even if the information is present at low levels, additional contrast resulting from structured illumination may help with detection. We can also note that for a low spatial frequency input, almost the whole pupil contributes to the OTF. However, the complex conjugation of one of the pupil function terms in Eq. 1 means that the phase terms almost cancel out. These low frequency components are thus only weakly sensitive to aberrations.

The IsoSense illumination method relies on the fact that if the sample is illuminated with a pattern containing several spatial frequencies (Fig. 4a, b), the spatial frequency content *F*(*r*) in the resulting fluorescence image would be *F*(*r*) = *I*(*r*) * *O*(*r*), a convolution of the frequency content of the illumination pattern *I*(*r*) and the frequency content of the object *O*(*r*). Therefore, at the pupil, multiple copies of the sample spatial frequency content would be created around the carrier frequencies that originated from illumination pattern (Fig. 4d). Appropriate selection of illumination pattern leads to a significantly improved sampling of the system OTF, especially at the high frequency periphery. As a practical compromise, to ensure sufficient pupil sampling we chose the pattern which permits two carrier frequencies to be visible diagonally in the image FFT, as depicted in (Fig. 4d). This is a simple but effective solution requiring no prior information about the sample.

**Figure 4:**
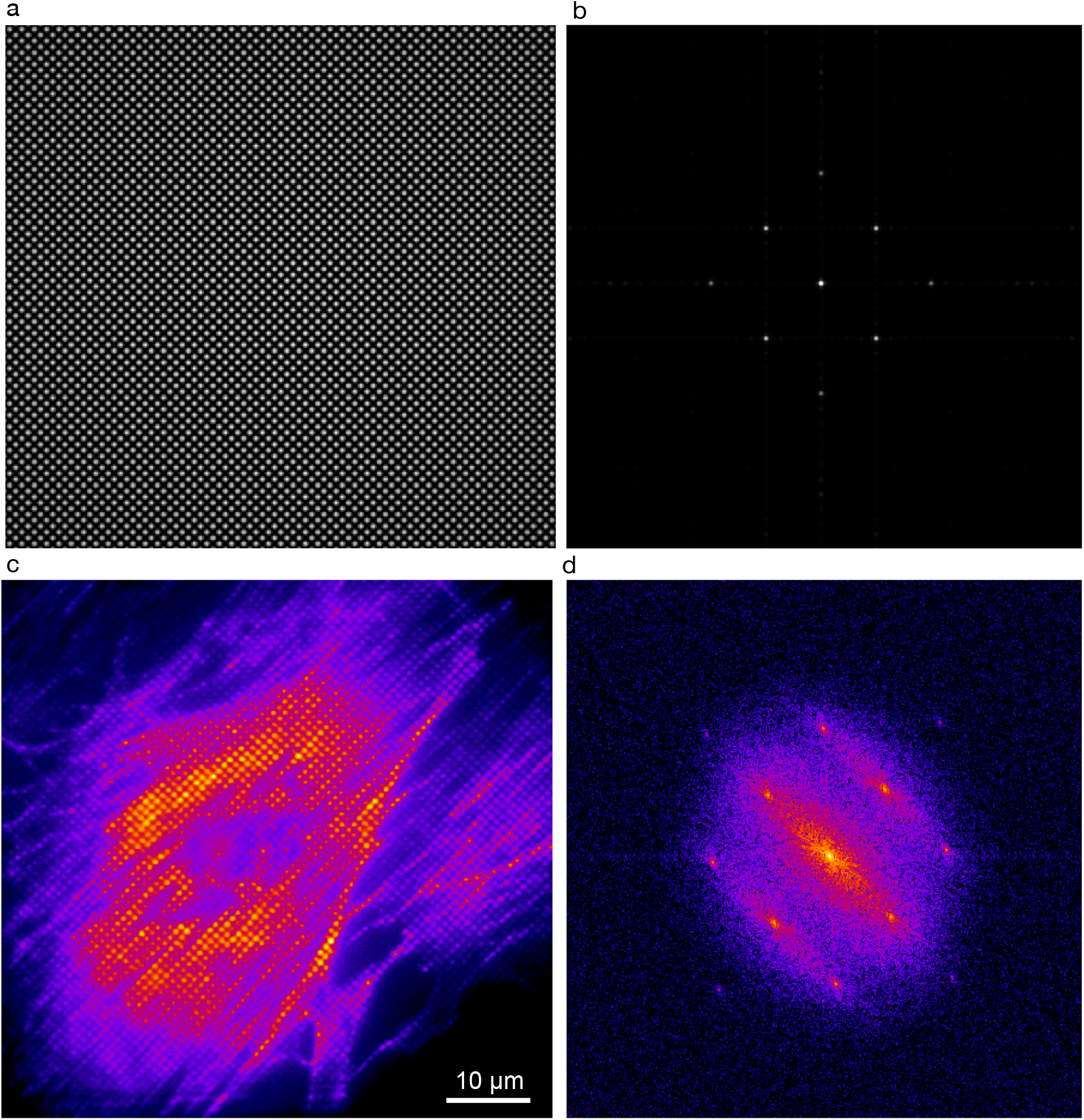
Structured illumination for reducing sample dependent effects on wavefront sensing accuracy. (a) simulated illumination pattern resulting from 4 beam interference. (b) Frequency content of the illumination pattern (c) an image acquired from a fixed bovine pulmonary artery endothelial cell showing actin (Alexa 488, phalloidin), illuminated with the sensing pattern. (d) Fourier transform of (c) demonstrates increased signal at higher spatial frequencies.

To generate custom illumination patterns, a grayscale phase mask *P*(*x*, *y*) was computed by multiplying two 2D sine waves (Eq. 2) and applied to the SLM (see section 4 for the description of optical set-up).

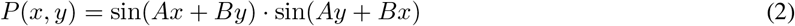

In the equation 2, *x*, *y* denote the vertical and horizontal coordinates in pixels, *A* controls carrier frequency and B the 2D pattern orientation. The resulting beams diffracted from the SLM were focused on the microscope pupil plane. A uniform regular interference pattern resembling an array of light beads was formed once all the beams were recombined at the sample plane after the objective lens. Figure 4a depicts a simulated illumination pattern resulting from interference of recombined beams. Figure 4b depicts a 2D Fourier transform of Fig. 4a and it clearly shows the main frequency components of the illumination pattern, that are in effect carrier frequencies for shifted copies of the sample structure information as shown in figures 4c and 4d of Alexa 488 phalloidin labelled actin in a fixed bovine pulmonary artery endothelial cell. (Fig. 4 c, d).

## 3 Interferometric wavefront sensing for deformable mirror control

A pre-requisite requirement for a reliable operation of sensorless AO is the ability to precisely control the adaptive element, in this case a deformable mirror (DM). The shape of the thin membrane forming the mirror can be controlled by applying the voltage to individual actuators that are situated behind the membrane that can push or pull at the corresponding membrane points. As there is no feedback on the position of the actuators or the shape of the membrane we measure the mirror shape with external wavefront sensing techniques. For DM control training and initial mirror flattening, we chose to use interferometric wavefront sensor-based approach [22].

An adapted version of Mach-Zehnder interferometer was built into the system to include a deformable mirror and part of the imaging system. The collimated and expanded beam from a He-Ne laser (R-33361, Research Electro-Optics) was split into object and reference paths and the reference path was diverted towards Interference camera using two steering mirrors M1 and M2, the steering mirrors were also used to introduce a wavefront tilt, necessary for single shot wavefront measurements. In the object path, immediately after the cube beamsplitter (BS013, Thorlabs), we have included one additional lens to ensure that at the DM pupil the light from He-Ne laser is collimated. The light was then reflected from the deformable mirror positioned at the conjugate pupil plane and reflected back from a flat silver mirror, positioned at the sample plane after the objective. The reflected light was picked up by 50:50 pellicle beamsplitter (BP150, Thorlabs) and launched towards the interference camera (XiQ MQ042MG-CM, Ximea), that was positioned at the conjugate pupil plane, where it interfered with the reference beam.

For wavefront sensing, the interferogram was processed with custom fringe analysis software using a LabView implementation. Fig. 5 illustrates major wavefront retrieval steps: The recorded interferogram (Fig. 5b) was transformed by an FFT (Fig. 5c) and the frequencies around the carrier were cropped, masked out and shifted back to the zero-frequency position (Fig. 5d). The inverse FFT was then computed. The phase profile corresponding to the optical phase of the wavefront was then obtained after phase unwrapping (Fig. 5e) using Goldstein 2D phase unwrapping algorithm [23]. Finally, Zernike modes were fitted to decompose the wavefront.

**Figure 5:**
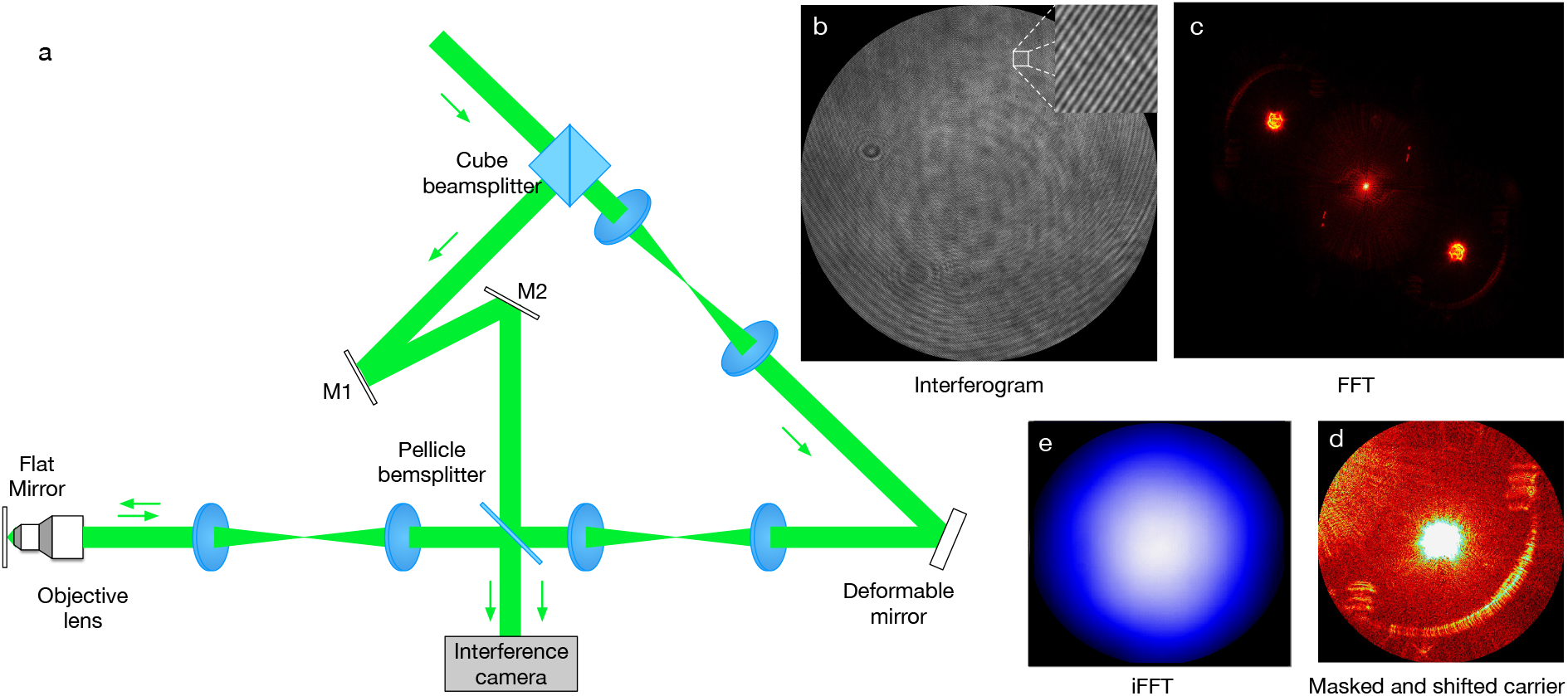
Interferometric wavefront sensing for mirror control. (a) a shematic showing the main components of the interferometer; (b) an iterferogram captured with the interference camera (c) FFT of (b); (d) masked and shifted high frequency carrier; (e) phase profile corresponding to the optical phase of the wavefront was then obtained after phase unwrapping.

The interferograms were captured and then used for (1) mirror training to produce Zernike modes and (2) mirror flattening. Mirror training was performed by capturing a set of interferograms obtained after poking individual actuators at various voltages (we used 5 steps at increments, each corresponding to 7.5% of full range). The wavefronts were decomposed into Zernike modes allowing the contribution of each actuator to a given Zernike mode to be calculated, giving a “poke matrix”. The control matrix is the inverse of this “poke matrix”. For a detailed description of the training method please see [24]. Once the control matrix was calculated, we have used the feedback from the interferometer to flatten the deformable mirror. This was typically carried out before the start of each cell-imaging experiment.

## 4 Optical set-up for demonstration of IsoSense in SIM

We chose a SIM set-up to demonstrate the IsoSense wavefront sensing method because the microscope already has a capability to produce arbitrary structured illumination patterns, that are necessary for wavefront sensing. We built an SLM-based SI microscope based on an optical light path design originating from John Sedat (UCSF), which will be published elsewhere in detail. A simplified schematic of the set-up is presented in figure 6.

**Figure 6:**
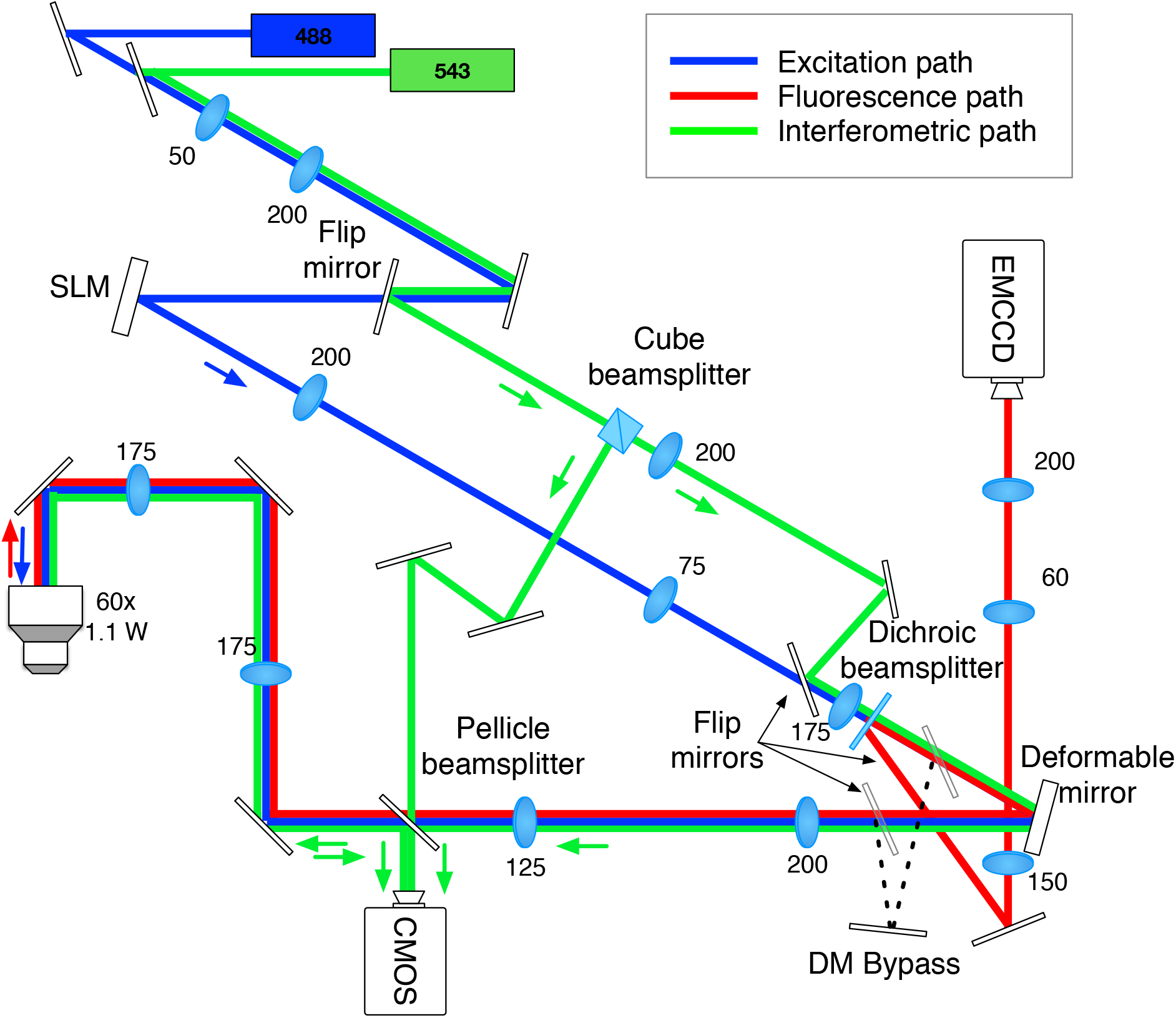
Optical setup for AO SIM imaging.

The light from two lasers: 488nm diode laser (LDM488.200.TA, Omicron Laserage), which is used for fluorescence excitation and 543 nm He-Ne (R-33361, Research Electro-Optics), which is used for interferometric wavefront sensing, were combined to a single path. The combined beams were launched at the SLM (HSPDM512-480-640, Boulder Nonlinear Systems, Inc.), which was positioned at the object plane conjugate. The diffracted beams from the SLM formed multiple spots on the deformable mirror (DM-69 BIL103, Alpao) which was positioned at the conjugate to the objective pupil plane. The DM was de-magnified to match the pupil size of the microscope objective (LUMFL N 60x/1.1W Olympus). The excitation beams interfered at the object plane to produce full-field structured illumination. Additional passive and active polarization control optics (not shown in the figure) were used to ensure optimal fringe contrast at the object.

The fluorescence light was collected by the objective lens and propagated back along the same path until the dichroic beamsplitter, where it was separated from the excitation path and launched towards an EMCCD camera (iXon Ultra 897, Andor).

The image timing and image acquisition were controlled using a custom Digital Signal Processing Unit (M62/67, Innovative Integration), driven by the Cockpit microscopy control environment that was developed in conjunction with UCSF. The SIM images were reconstructed using the fairSIM [25] plugin for ImageJ.

## 5 Results and Discussion

### 5.1 Sample preparation

FluoCells prepared slide #1 (F36924) containing bovine pulmonary artery endothelial cells stained with red-fluorescent MitoTracker Red CMXRos, green-fluorescent Alexa Fluor 488 phalloidin (actin), and DAPI was purchased from Invitrogen.

*Drosophila* macrophages (hemocytes) were isolated from wandering third-instar larva expressing Jupiter:GFP and Histone:RFP ubiquitously, as described in [26]. To prepare samples for imaging, 4-5 larvae were collected, washed with gentle vortexing and the cuticle pierced to release the hemolymph in 200l Schneiders culture medium (Thermofisher) in a 35 mm glass bottom dish (MatTek). The preparation was allowed to stand in a humid chamber for at least 30 minutes to allow macrophages to adhere before additional medium was added for imaging. This preparation yielded predominantly plasmatocyes with occasional larger, flatter lamellocytes [27].

### 5.2 Imaging and adaptive optics correction

Custom illumination patterns have long been used in SIM to down-shift high spatial-frequency components in order that they fit within the pass band of the system. However, using SIM reconstructions for image quality based adaptive wavefront sensing is impractical for two reasons: (1) a single reconstruction requires multiple images to be taken, increasing the effects related to phototoxicity and bleaching, (2) if the microscope OTF is compromised by severe optical aberrations, common reconstructions would fail [28]. Furthermore, as the IsoSense method relies on indirect estimates of OTF quality through the high spatial frequency components of raw images, no additional reconstructions are needed. The image quality metric representing system performance in SIM mode can be calculated from a single image.

IsoSense is particularly well suited for SIM imaging systems that rely on dynamic optical components for creating SIM patterns, as the same illumination control element can be used for both: SIM imaging and wavefront sensing. Therefore, we chose a liquid crystal SLM based SI microscope for initial proof of concept demonstration to verify the performance of the method.

Prior to imaging we trained the mirror and flattened the system using interferometer feedback for Zernike modes 5-100, following the Noll notation [29]. To get the reference to the “system flat” correction and to correct non-common path aberrations we performed image-based correction for sub-diffraction sized fluorescent beads (0.1 micrometer diameter) scattered on the coverslip, using distilled water immersion. Once the system calibration is completed the resulting control matrix and “system flat” file can be used for further imaging experiments if the optical system remains unchanged. Prior to each individual imaging experiment, a sensorless adaptive correction was performed for each isoplanatic patch in the object to correct for sample -induced aberrations. “System flat” correction was determined for the sake of having a fair comparison between AO-corrected and images for the equivalent non-AO microscope where the deformable mirror was bypassed as illustrated in Fig. 7. Therefore “system flat” includes both interferometric correction and sensorless correction of non-aberrating sample to account for other system aberrations. For system flat correction we corrected first 5-22 Zernike modes in two iterations. “System flat” correction using beads is not necessary for day to day imaging experiments, as other system aberrations are also detected and corrected during sensorless AO correction of sample induced aberrations.

**Figure 7:**
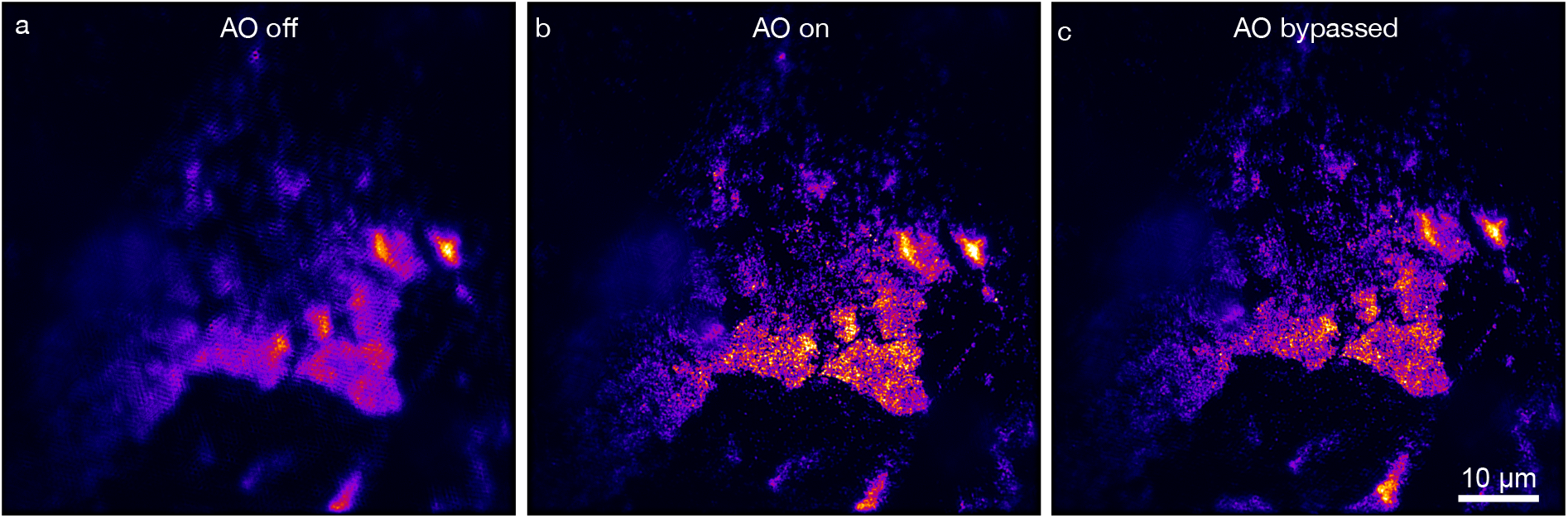
SIM reconstruction of 2D bead clusters on the coverslip represent a non-aberrating object. In a well-aligned system “system flat” AO correction setting (b) should provide similar image quality as when AO is bypassed (c). When the mirror is switched off (a), the membrane relaxes, significantly affecting image quality.

Next, we corrected sample and non-common-path aberrations. Sample induced aberrations are caused by the sample structure and immersion medium, while non-common path aberrations can occur due to the slightly different optical paths used during the system calibration (interferometric path) and imaging (excitation path, fluorescence path) shown in Fig. 6. From prior experiments, we knew that the first two orders of spherical aberration tend to dominate due to refractive index mismatch between sample and immersion medium. Also, we used Schneiders *Drosophila* medium (ThermoFisher) with a water immersion objective, therefore the immersion medium was not fully compatible with the immersion lens. After correcting two orders of spherical aberration, which were present due to refractive index mismatch, we have corrected coma, astigmatism and trefoil – these aberrations most likely were non-common path aberrations that were caused by the presence of pellicle beam splitter in the interferometric wavefront sensing path during interferometric mirror flattening. For cell images presented in our paper, correcting Zernike modes (5-10, 22) was sufficient, however, this might be different for different types of cells and tissues. To ensure robust noise performance during sensorless AO correction we were using 5 images per mode.

The first example, a *Drosophila* larval macrophage expressing Jupiter::GFP labelling microtubules, demonstrates the effect of AO correction in SIM (Fig. 8). The effect of correction is much less obvious in corrected (Fig. 8b) and “system flat” (Fig. 8a) widefield images, however, it is much more prominent in SIM reconstructions where AO correction results in reduced SIM reconstruction artefacts (Fig. 8d) compared to the case where the system flat was used (Fig. 8c).

**Figure 8:**
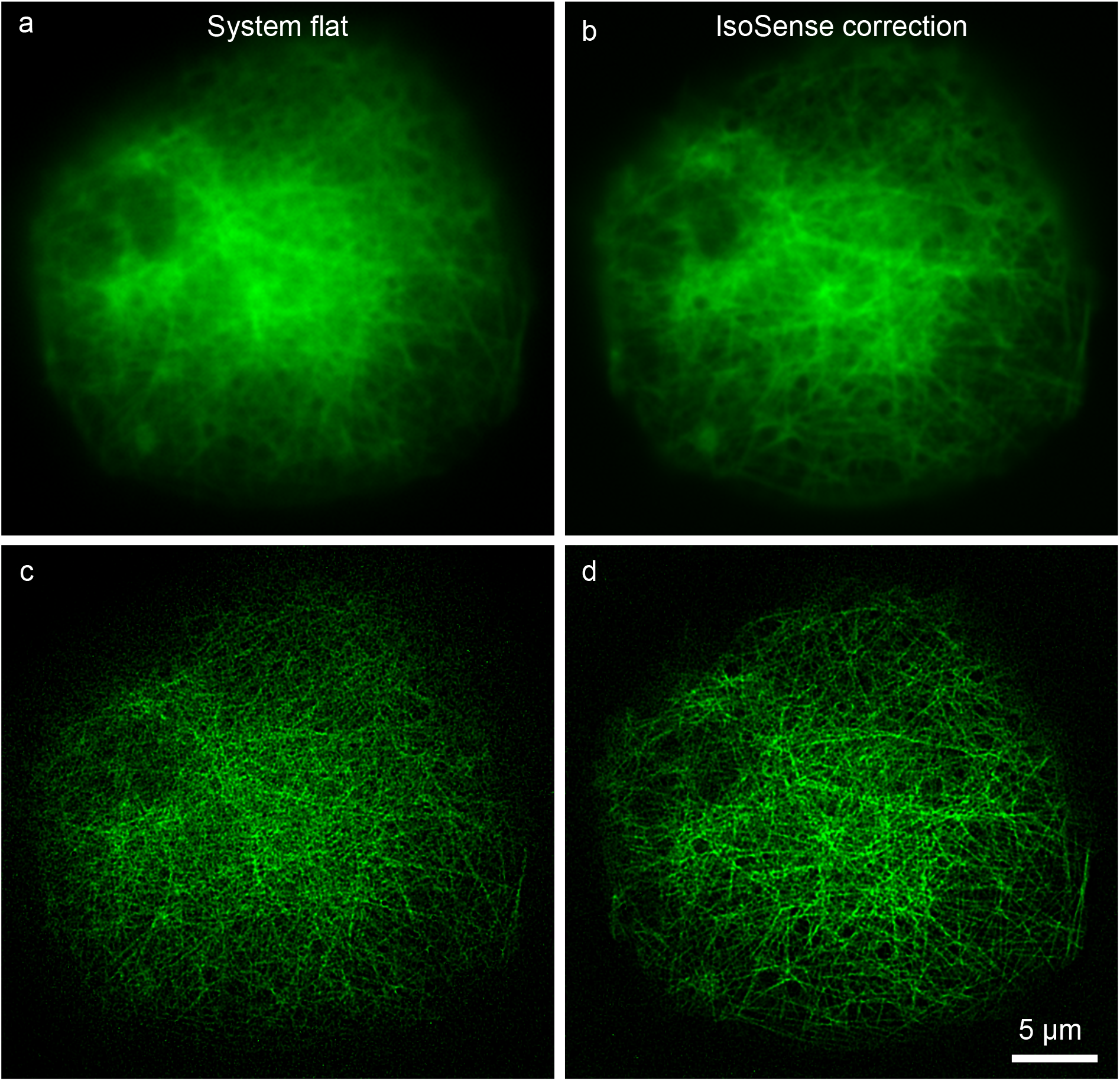
AO correction for live SIM imaging of a macrophage expressing Jupiter::GFP (labelling microtubules). IsoSense corrected images (b) and (d) contain fewer artefacts than the images (a) and (c) that were obtained using system flat correction. The benefit of correction is much more apparent in SIM reconstructions (c) versus (d) (bottom row) than in pseudo-widefield reconstructions (a) versus (b) (top row).

The second example clearly demonstrates the benefit of the IsoSense wavefront sensing approach. Here the aberrations were first corrected with sensorless AO using wide-field illumination (Fig. 9 a, c) and then the correction was repeated using the IsoSense approach. Fig. 9 represents a typical example where the majority of object structure has a clear preferential orientation, resulting in insufficient sampling of the OTF. As a result, a better correction was achieved along the direction that is perpendicular to the orientation of the majority of microtubules; along the other direction the microtubules appear washed out. Again, this is particularly visible in the SIM reconstruction (Fig. 9c). IsoSense approach led to better overall correction, preserving all the structures in the sample (Fig. 9d).

**Figure 9:**
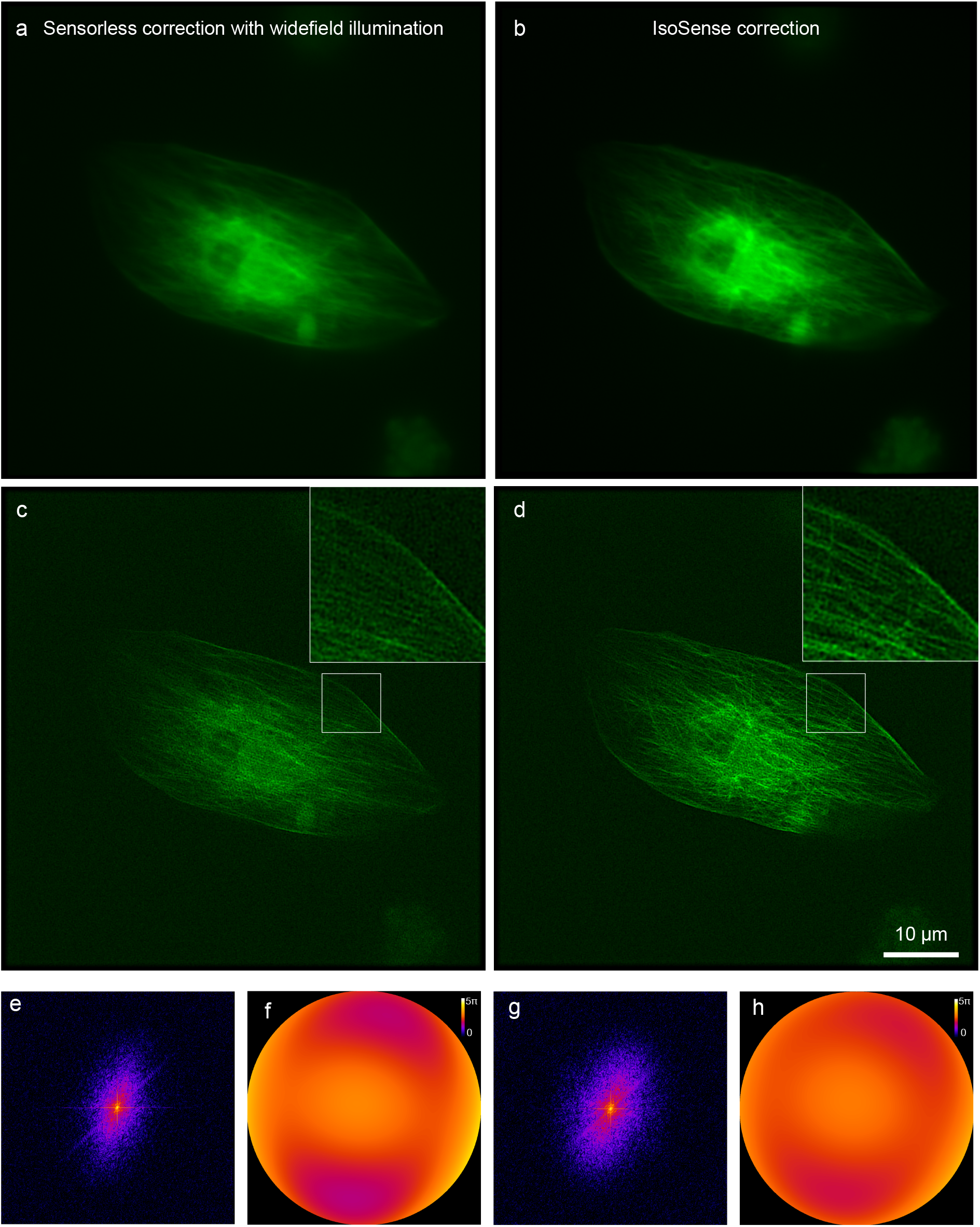
AO corrections for cellular imaging with SIM. (a) and (c) represent widefield and reconstructed images of Jupiter::GFP expressing lamellocytes labelling microtubules, where image quality based wavefront sensing was performed with widefield illumination. In (b) and (d) custom illumination patterns for imaging quality sensing were used. (c) demonstrates higher quality correction along the direction corresponding with preferential fiber orientation, while (d) shows more isotropic and better overall correction. (e, g) are the amplitude components of Fourier transforms of the widefield images and (f, h) the detected phase profiles used for correction (in radians).

## 6 Conclusion

The IsoSense structured illumination based sensorless AO approach removes the barriers to accurate image quality metric calculation, commonly caused by anisotropic object structure or lack of sharp and well-defined structures. IsoSense improves the sampling of the microscope OTF by filling it with spatial frequency-shifted image information the in otherwise poorly populated OTF regions. Consequently, it allows us to detect the aberration components in the corresponding pupil regions. Finally, we have demonstrated the performance of our method in a SIM set-up for imaging live *Drosophila* macrophage cells expressing a GFP tagged microtubule marker, where our method effectively improved the accuracy of AO correction and image quality.

This demonstration used a versatile SLM to generate the patterns. However, the method should be compatible with alternative pattern generation techniques that do not require an active optical element (speckle patterns, a phase mask, transmissive masks). Even though in this paper the IsoSense method was demonstrated using interference patterns, coherent illumination is not inherently required. The method can be easily extended to compatible imaging systems with low coherence or incoherent illumination, such as light sheet structured illumination microscopy [30] or lattice light-sheet microscopy [31].

The technique can also be extended to include adaptive pattern generation. For example, when imaging strongly aberrated objects, the carrier frequencies could be positioned at lower frequencies, closer to pupil center, in the first iteration and gradually shifted to higher frequencies during subsequent correction iterations. Alternatively, the pattern generation algorithm can be automated to identify inadequately sampled OTF regions and adjust the phase mask to generate the carrier frequencies that fill these blind spots.

We anticipate that IsoSense will be useful in conjunction with other imaging techniques that rely on adaptive optics for aberration correction. AO light-sheet microscopy [32] is particularly well suited to benefit from IsoSense as these techniques already contain both AO and excitation pattern modulation device.

## Acknowledgments

We thank John Sedat for fruitful discussions and for sharing his optical light path design for a SLM-based SIM system with adaptive optics, on which we based our microscope design.

The datasets generated during and/or analysed during the current study are available from the corresponding author on reasonable request.

